# Estrogen accelerates heart regeneration by promoting inflammatory responses in zebrafish

**DOI:** 10.1101/616250

**Authors:** Shisan Xu, Fangjing Xie, Samane Fallah, Fatemeh Babaei, Lina Zhu, Kin Fung Wong, Yimin Liang, Rajkumar Ramalingam, Lei Sun, Xin Wang, Yun Wah Lam, Shuk Han Cheng

## Abstract

Sexual differences are observed in the onset and prognosis of human cardiovascular diseases, but the underlying mechanisms are not clear. Here, we report that zebrafish heart regeneration is faster in females, can be accelerated by estrogen and suppressed by estrogen-antagonist tamoxifen. Injuries to the heart, but not other tissues, increased plasma estrogen level and expression of estrogen receptors, especially *esr2a*, in zebrafish hearts. The resulting endocrine disruption induces the expression of female-specific protein vitellogenin in male zebrafish. Transcriptomic analyses suggested heart injuries triggered more pronounced immune and inflammatory responses in females. These responses, previously shown to enhance heart regeneration, could be enhanced by estrogen treatment in males and reduced by tamoxifen in female. Furthermore, a brief exposure to estrogen could precondition zebrafish for an accelerated heart regeneration. Altogether, this study reveals that heart regeneration is modulated by an estrogen-inducible inflammatory response to heart injury. These findings elucidate a previously unknown layer of control in zebrafish heart regeneration and provides a new model system for the study of sexual differences in human cardiac repair.

## Introductiono

Cardiovascular diseases (CVD) are the primary cause of death worldwide: causing the death of 17.9 million people in 2015, more than 30% of the global mortality (Roth et al., 2017). Gender differences have been reported on risk factors, clinical manifestation and recovery of CVD (reviewed by Maas and Appelman, 2010; Ostadal and Ostadal 2014; EUGenMed et al., 2015). In various mammalian models of cardiac defects, females consistently demonstrate a lower mortality, less severe disease phenotypes and better functional recovery than the male counterparts (Du 2004; Czubryt et al., 2006; Du et al., 2006). An understanding of the molecular mechanism underlying these gender differences will provide an important gateway towards better CVD prevention and treatment. While the protective effects of estrogen on the cardiovascular system have been widely reported, most of these studies focused on how estrogen ameliorates CVD risk factors, by the maintenance of vasodilation and in the prevention of atherosclerosis (Pare et al., 2002; Mendelsohn 2002; Babiker et al., 2002; Fliegner et al., 2010). But estrogen may play more direct roles in cardiac development, functions and pathology, as estrogen can protect cardiomyocytes from apoptosis caused by ischaemia–reperfusion in vitro (Liu et al., 2011). Gene expression profiles of mammalian cardiomyocytes (CM) are sexually dimorphic (Isensee et al, 2008, Witt et al, 2008, Tsuji et al, 2017), and estrogen receptor expression is deregulated in some cardiomyopathies (Mahmoodzadeh et al., 2006). Despite the apparent involvement of estrogen in CVD, clinical translation of these findings has been lagging, partly due to inconclusive results from large-scale studies on the benefits of post-menopausal hormone replacement treatment on CVD prevention (Low et al., 2002). This is attributed, at least in part, to the lack of a mechanistic understanding of the roles of estrogen in cardiomyocyte biology, which prevents optimal study design and subject selection in clinical studies (Whayne & Mukherjee, 2015).

While adult human cardiomyocytes are virtually unable to re-enter the cell cycle, other vertebrates, including zebrafish, newts and axolotls, demonstrate a remarkable ability to regenerate their hearts (reviewed by Garcia-Gonzalez and Morrison 2014; Vivien et al., 2016). Among these organisms, zebrafish has emerged as one of the most well-established model organisms for the study of heart regeneration. Even after the resection of up to 20% of its ventricle, a zebrafish can completely regenerate its heart within 2 months with no apparent scarring (Poss et al., 2002). Apart from physical amputation, cryoinjury (Chablais et al., 2011; González-Rosa et al., 2011) and genetic ablation (Wang et al., 2011) of cardiomyocytes can also trigger proliferation, usually measured by the expression of proliferating cell nuclear antigen (PCNA) or phosphor-histone H3. Heart regeneration is not only limited to the proximity of the wound, but involves the precise coordination of cardiomyocytes proliferation, migration (Itou et al., 2012) and extracellular matrix remodelling (Wang et al., 2013) throughout the heart. Fate mapping experiments have established that the proliferating cardiomyocytes originated from mature cardiomyocytes (Jopling et al., 2010; Senyo et al., 2013; Zhang et al., 2013). The rapid disappearance of the sarcomeric structures and re-expression of embryonic genes, such as embryonic cardiac myosin heavy chain gene (embCMHC), suggest that regeneration largely involves the expansion of dedifferentiated cardiomyocytes (Jopling et al., 2010; Sallin et al., 2015). Biochemical and gene expression analyses have identified the involvement of growth factors such as *tgfb1, pdgf, shh* and *igf2b*, and signalling pathways such as Notch, Jak1/Stat3 and NFκB in zebrafish heart regeneration (reviewed by Kikuchi et al., 2014; González□Rosa et al., 2017). A recent study has attributed the remarkable regenerative capacity of zebrafish heart to this organism’s enhanced inflammatory and immune responses, in comparison with other fish species (Lai et al., 2017). Hypoxia, as a result of cardiac damage, also plays a positive role in cardiomyocyte proliferation (Jopling et al., 2012), whereas hyperoxia and miR-133 inhibit regeneration (Yin et al., 2012).

While the zebrafish heart is different from the human counterpart on cellular and anatomical levels (Hu et al., 2000; Jensen et al., 2013), the conservation of the genetic circuitry between zebrafish and mammals (Howe et al., 2013), and the technical and genetic resources offered by the zebrafish research community, make this organism an ideal model for the study of the gender bias associated with human CVD. Some observations have suggested a role of sex hormones in zebrafish cardiac development and functions. For instance, inhibitors of estrogen synthesis induced phenotypes similar to congestive heart failure and tamponade in zebrafish embryos (Allgood et al., 2013). These conditions could be reversed by estradiol (E2) treatment, which has also been shown to affect heart rates during zebrafish embryonic development (Romano et al., 2017). Despite these findings, the entire literature on zebrafish heart regeneration is based on studies on a single sex (usually male) and sexual differences in regeneration have never been investigated. In this study, we examine, for the first time, the sexual dimorphism of zebrafish in heart regeneration and in response to factors that influence the rate of cardiac regeneration.

## Results

### Zebrafish heart regeneration is sexually dimorphic

Female and male zebrafish, matched for age and weight, were subjected to cardiac damage by cryoinjury. On day 7 post cryoinjury (7 dpc), female hearts contained a significantly higher number of PCNA-positive cells (Fig 1A) and less vimentin immunoreactivity (Fig 1C) than in the male counterpart (Fig 1B and 1D), indicating more cell proliferation and fewer scar-forming fibroblasts in the regenerating female heart. After one month of regeneration, female hearts contained significantly smaller scar tissues compared to males (Fig 1E, F). Uninjured female hearts contained a significantly higher number of PCNA-positive cells than the males (Fig. 1A, B), indicating a higher baseline proliferative activity in female cardiomyocytes. Inflammation is known to be essential for the initiation of regenerative responses (Kyritsis et al., 2012). Our data showed that the number of L-plastin positive leukocyte was higher in the injured area of female fish than that in male fish at 1 dpc (Fig. S1). We compared the relative scar volume in female and male hearts, and showed that while the scar volume was similar in both sexes at 1 dpc, the scar volume in male fish was about two folds larger than in female fish at 30 dpc (Fig 1 E, F). Together, these data support the sexual dimorphism of zebrafish heart regeneration, with females regenerating their hearts faster than males.

**Figure 1.**
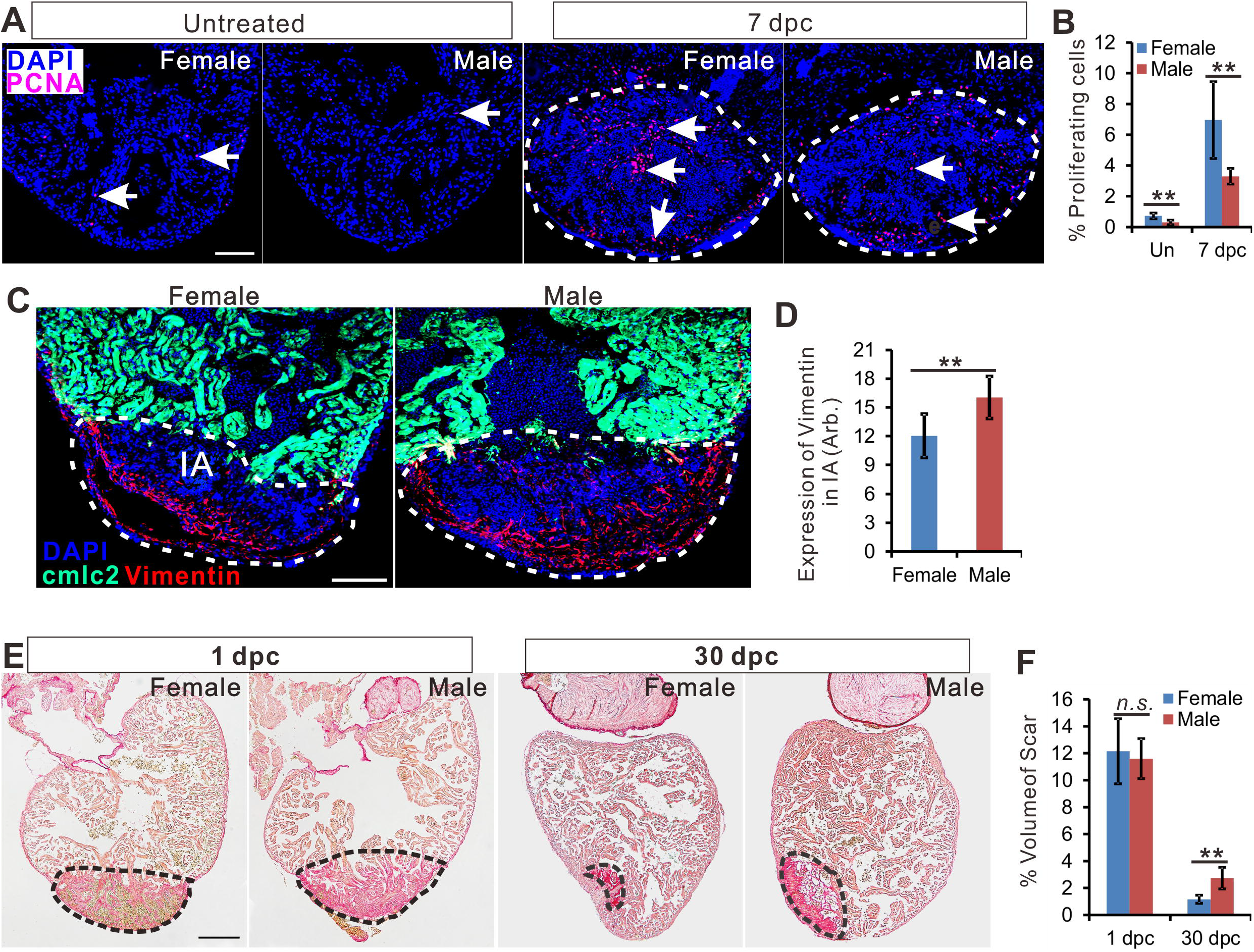
Zebrafish heart regeneration is sexually dimorphic. **A.** PCNA immunofluorescence (red) in the heart of untreated female, untreated male, 7dpc female (c) and 7 dpc male. **B**. Quantification of percentage of PCNA positive cells (mean±SD, n=7) in panel A. **C**. Vimentin immunofluorescence (red) in female and male Tg (cmlc2: eGFP, green) zebrafish hearts at 7 dpc. **D**. Quantitation of vimentin expression in the injured area of female and male fish (marked by dashed lines, mean±SD, n=8). **E**. Picrosirus red staining of female and male zebrafish hearts at 1 dpc and 30 dpc. **F**. Quantitation of scar volume (marked by dashed lines, mean±SD, n=9~12) between females and male fish. Scale bars: 200μm. Two-tail t-test, **p<0.01. Un: untreated, Dpc: days post-cryoinjury, CI: Cryoinjury.

### Endocrine disruption of zebrafish following cardiac injury

Our observations on the sexual dimorphism of zebrafish heart regeneration prompted us to examine the involvement of estrogen in this process. Towards this goal, the expression of all three nuclear ERs (Lu et al., 2017) was examined in zebrafish hearts after cryoinjury or sham operation, as well as in the heart of injured fish (untreated). Sham operation was performed by the identical procedures that preceded cryoinjury, including the opening of chest skin and tissue to expose the heart. Fig. 2A shows that at 7 days post-surgery, the expression of *esr1* and *esr2a* was significantly induced (~3.5-fold for *esr1* and ~9-fold for *esr2a*) in female hearts. Interestingly, the expression of these two ERs in male hearts was also significantly increased, though their increase was not as pronounced as in female heart (~2-fold for *esr1* and *esr2a*, Fig 2B). Sham operation also induced the expression level of *esr2a* (in female) and *esr1* (in both sexes), but to a lesser extent compared to the effect of cryoinjury. *Esr2b* was suppressed by both cryoinjury and sham operation in either sex.

**Figure 2.**
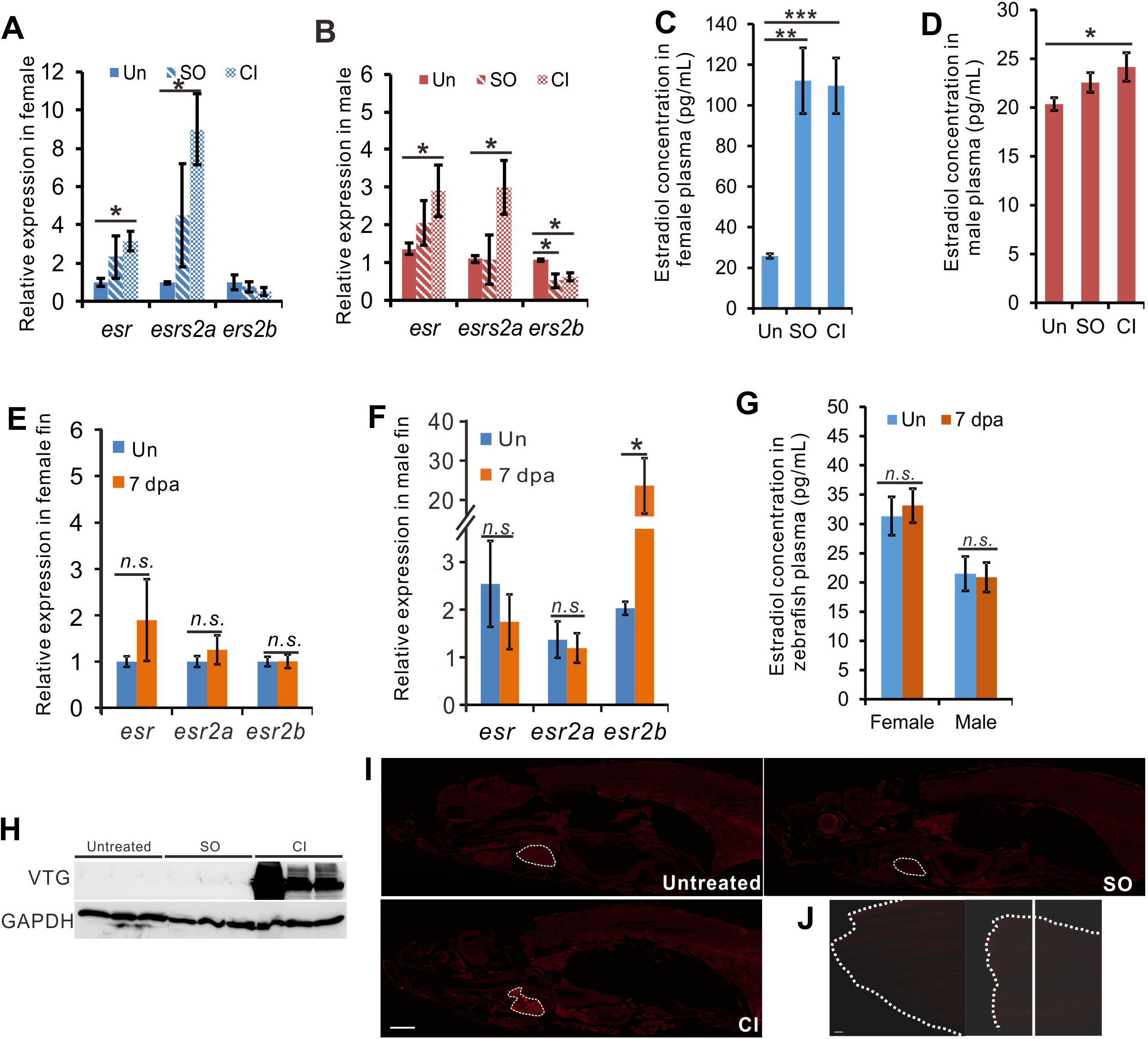
Cardiac damage induces feminisation of zebrafish. **A-G**. q-PCR to show the expression of estrogen receptor genes in the heart (A, B) and fin (E, F) of zebrafish 7 days after CI; and plasma E2 concentration in zebrafish with heart (C, D) or fin (G) injury at 7 days. N=3, two-tail t test (A, B, E, F, G), and one-way ANOVA with LSD post hoc test (C, D), *p<0.05, **p<0.01, n.s, no significant. **H.** Detection of vitellogenin by Western blotting in plasma collected from untreated male zebrafish and fish one day after SO and CI. **I**, Detection of VTG (red) in an untreated male zebrafish and in male zebrafish on day 7 after heart CI and SO. Scale bars: 1mm. **J**. Detection of VTG in uninjured caudal fin and on 7 days after fin amputation. The dashed white line showed the shape of fin, and the white line showed the amputation site. Bars: 1mm. Un, untreated, SO, sham-operation, and CI, cryoinjury.

As ERs are estrogen inducible genes, we asked whether the level of E2 in those fish was also affected after heart injury. Consistent with the upregulation of *esr1* and *esr2a*, the plasma E2 level was increased by ~5-fold in sham-operated and cryoinjured female fish (Fig. 2C). Plasma E2 in male fish was also observed to increase by 20% and 10% after cryoinjury and sham-operation respectively (Fig. 2D). However, the expression of *esr1* and *esr2a* in the fin was not significantly altered after amputation in either female or male fish (Fig 2E, F), and the plasma E2 levels after caudal fin amputation were virtually unchanged as compared to untreated controls (Fig. 2G). Overall, these data suggest that injury to the heart, but not other tissues, induces estrogen secretion and response in zebrafish.

To test the biochemical consequence of this hormonal change, we utilized the sexual dimorphism in zebrafish plasma proteins (Babaei et al., 2013; Li et al., 2016). Quantitative proteomic analysis was used to study the relative abundances of 18 known sexually dimorphic plasma proteins in the plasma collected from untreated, sham-operated and ventricular amputated male zebrafish (Fig. S2A and Table S1). Most female-biased plasma proteins increased after cardiac damage, as compared to sham operation, except Nucleoside Diphosphate Kinase (NDPK), a protein known to be marginally female-biased (Li et al., 2016). Conversely, all but two male-biased plasma proteins were downregulated after ventricular amputation. As the two outliers, Myoglobin and Phosphoglycerate Kinase 1, are expressed predominately in muscle, the increase of these proteins in plasma might be a result of the release of cardiomyocyte content into circulation. These data indicate that although the impact of heart injury on estrogen level in males was relatively minor, it was sufficient to shift the gender characteristic of the male zebrafish plasma towards a more feminised composition.

Our plasma proteomic analysis indicates the presence of vitellogenin isoforms in the plasma of male zebrafish after heart injury (Fig. S2B). Western blotting detected vitellogenin in the plasma of male zebrafish as early as day 1 after both cryoinjury (Fig. 2H) and heart amputation (data not shown), as compared to sham or uninjured fish. Vitellogenin is specifically expressed in females (reviewed by Hara et al., 2016) and are routinely used as biomarkers for endocrine disruption in males (García-Reyero et al., 2004; Scott and Robinson, 2008). As vitellogenin has been reported as an acute phrase response protein (Tong et al., 2010; Zhang et al., 2011), it is possible that vitellogenin was induced in cardiac-damaged males as a result of infection. However, the abundance of vitellogenin in the plasma was much higher in heart-damaged than in sham-operated fish, even though the latter also suffered from extensive tissue damages. Moreover, plasma proteomics showed that most of the known acute phase response proteins (Roy et al., 2017), such as the complement and coagulating factors, were down-regulated in the heart regenerating fish as compared to the sham (Fig. S2B and Table S2). This suggests that the presence of vitellogenin in male plasma after heart injury is not a consequence of injury-related infection or inflammation; but is specifically associated with heart regeneration. We then examined the tissue distribution of vitellogenin in male zebrafish during heart regeneration. Whole mount immunohistochemistry showed that vitellogenin accumulated in the injured heart of male fish (Fig. 2I) and was not observed in the proximity of the chest wound of the sham-operated fish. This indicates that the vitellogenin accumulation in male fish is specifically associated with cardiac damages and is not related to general wound healing. Moreover, vitellogenin was not undetectable on the male caudal fin on day 7 after amputation (Fig. 2J), confirming that vitellogenin accumulation is not a general consequence of tissue regeneration or repair. And individual tissue immunohistochemistry analysis confirmed the result that vitellogenin was detected in the heart, but not in the gill, kidney and liver (Fig. S2C). Vitellogenin was observed in the entire regenerating heart, not restricted to the wound.

### Estrogen promotes heart regeneration in zebrafish

What is the effect of estrogen on heart regeneration? At 7 dpc, tamoxifen inhibited the expression of *esr2a* in female heart (Fig. 3A), while E2 increased the expression of *esr2a* in male heart (Fig. 3B). Male zebrafish exposed to E2 after cryoinjury displayed a ~4-fold increase in cardiomyocyte proliferation in the vicinity of the injured area (Fig. 3C, D). E2 treatment of male fish also increased cardiomyocyte dedifferentiation, as judged by the expression of embCMHC (Fig. 3E-G). Conversely, treatment of female fish with tamoxifen, an antagonist of estrogen receptors (Jordan, 2006; Xia et al., 2016), resulted in a ~4-fold decrease in cardiomyocyte proliferation in the vicinity of injured area (Fig. 3A). and ~2-fold decrease in embCMHC expression (Fig. 3E-G). Furthermore, E2 treatment of male fish increased the level of vitellogenin in the regenerating hearts, while tamoxifen treatment of female fish reduced this accumulation (Fig. 3H-J).

**Figure 3.**
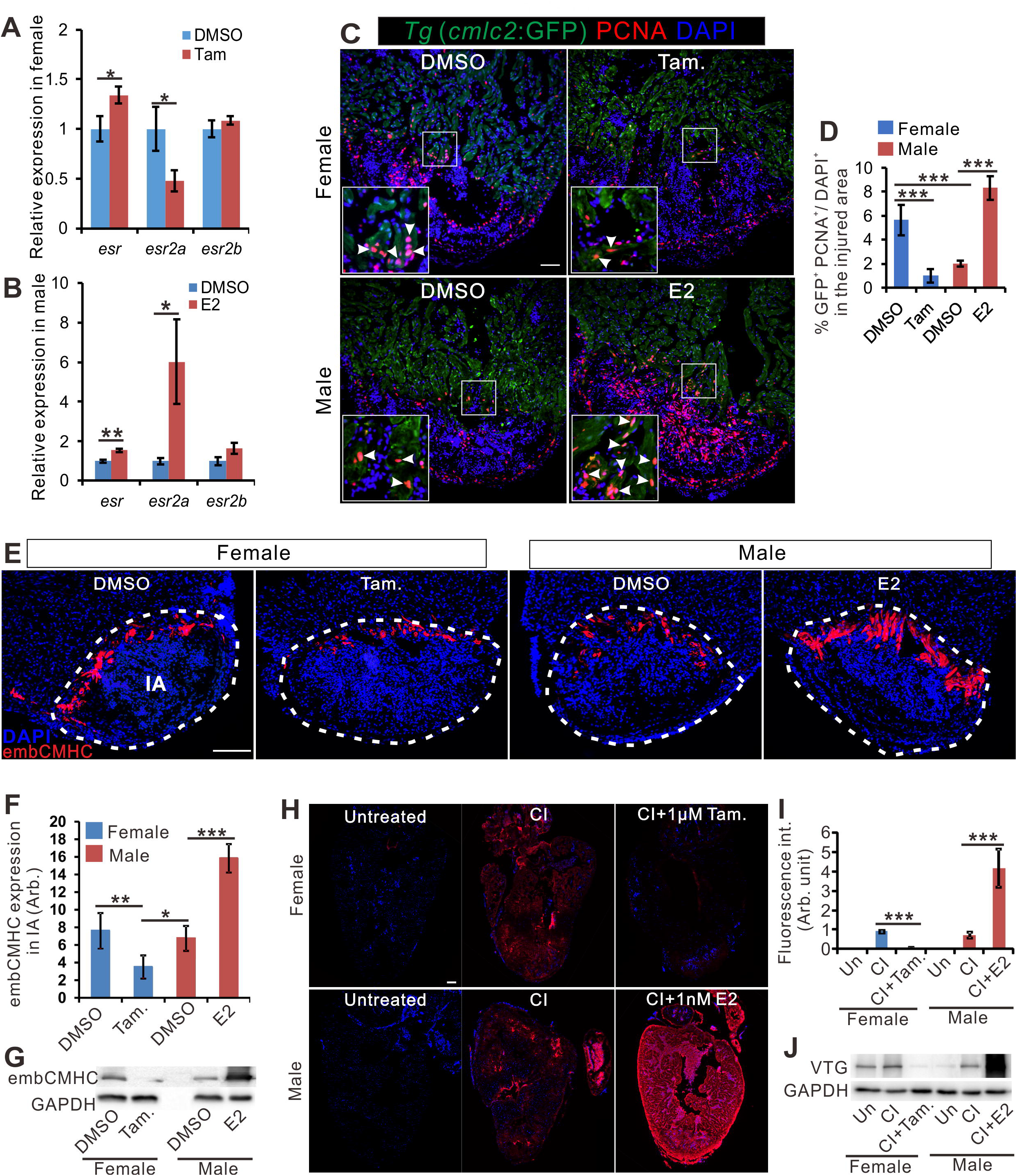
Estrogen promotes regenerative programme in zebrafish heart. **A-B**. q-PCR to show the expression of estrogen receptor genes in the heart of female fish (A) and male fish (B) with DMSO, tamoxifen or E2 treatment at 7 days after cryoinjury. **C**. PCNA immunofluorescence (red) in the heart of male *Tg* (*cmlc2*: eGFP) zebrafish exposed to 1nM E2 for 7 days after CI. Scale bar: 100μm. **D**. Quantification of proliferating cardiomyocytes in panel A (mean±SD, n=5) in injured areas. **E**. embCMHC immunofluorescence (red) in the heart of untreated female, female treated with 1μM tamoxifen (Tam.), untreated male, and male treated with 1nM E2 for 7 days after CI. Scale bars: 200μm. **F**. Quantification of embCMHC staining in panel C (mean±SD, n=4~5) in injured areas. **G**. Expression of embCMHC in the samples shown in C, as detected by Western blotting. One-way ANOVA test, *p=0.05, **p=0.01, ***p<0.001. **H**. Vitellogenin (VTG) immunofluorescence (red) in the heart of untreated female, 7 dpc female, 7 dpc female treated with 1μM Tamoxifen, untreated male, 7 dpc male, and 7 dpc male treated with 1nM E2. **I**. Quantification of VTG staining (mean±SD, n=4~5) in panel A. Scale bar: 100μm. One-way ANOVA with LSD post hoc test, ***p<0.001. **J**. Expression of VTG in the samples shown in panel A, as detected by Western blotting. Un, untreated, SO, sham-operation, and CI, cryoinjury.

We then investigated whether tamoxifen and estrogen affect the scar degradation after heart injury. Tamoxifen significantly increased the scar volume in female fish (Fig. 4A, B), whereas E2 speeded up scar degradation in both female and male fish. Although E2 treatment could also accelerate heart regeneration in females, the effect was less pronounced (~2-fold reduction of scar size) than in males (~3-fold reduction of scar size, Fig. 4A, B). Moreover, echocardiographic measurements, that allowed the non-invasive measurement of cardiac performance during regeneration (Wang et al., 2017), indicated that E2 treatment of male fish after cryoinjury accelerated the restoration of Fractional Shortening (FS) time to the pre-injury level (Fig. 4C). FS is a parameter that indicates the cardiac contractile force and has previously been shown to correlate with zebrafish cardiac recovery during regeneration (Hein et al., 2015). The increase in FS recovery by E2 treatment confirms the promoting effect of this hormone on the recovery of physiological functions after cardiac damage in male zebrafish. Overall, our data showed that estrogen promoted the regenerative programmes and the recovery of heart function.

**Figure 4.**
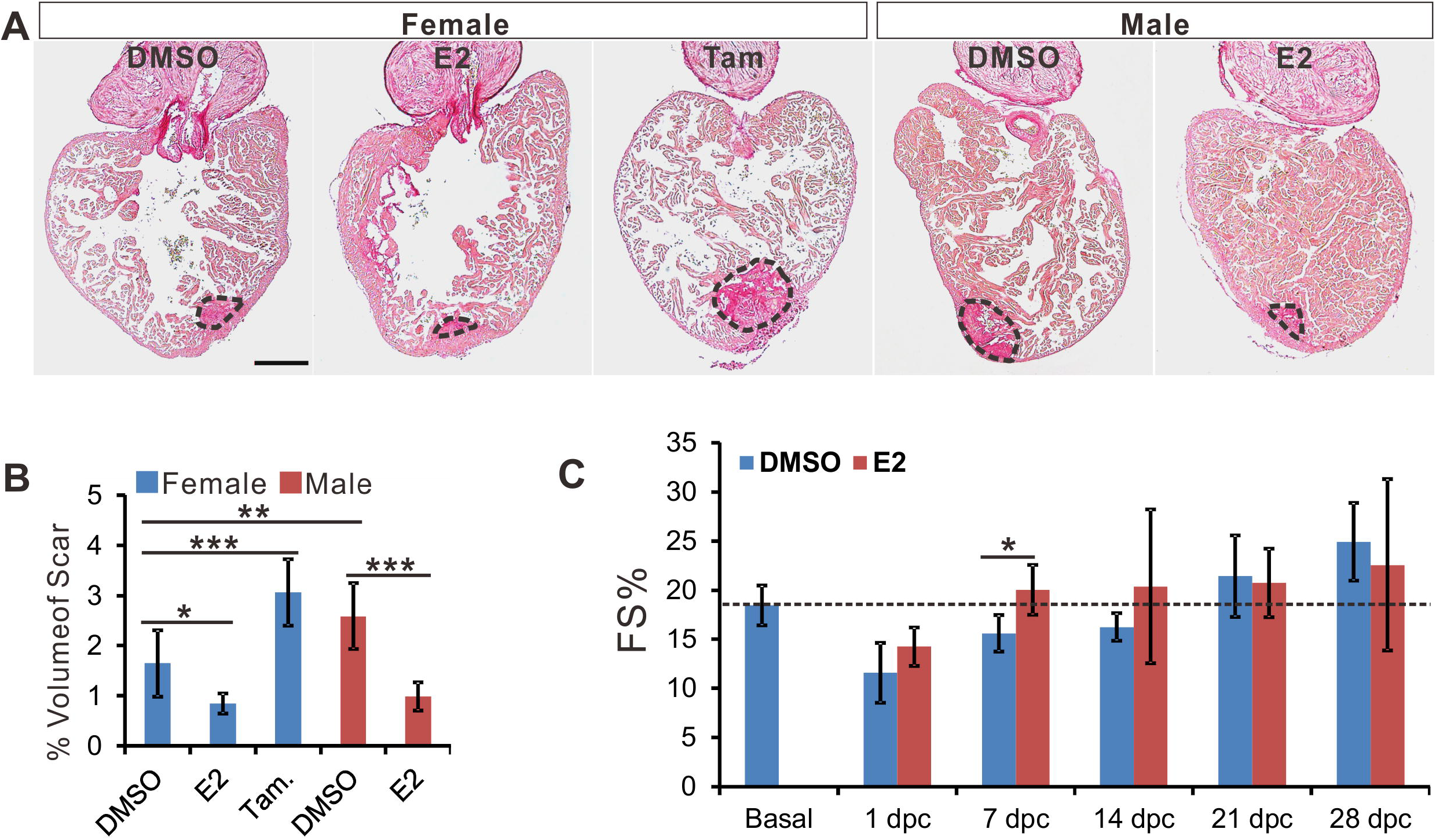
Estrogen accelerates scar reduction and promotes the recovery of cardiac functions. **A**. Picrosirus red staining of the heart from female zebrafish treated with DMSO, E2 (1 nM) and Tamoxifen (1 μM); and male zebrafish treated with DMSO and E2 (1 nM) at 30 dpc. Scale bar: 200μm. **B.** Quantitation of scar volume of the samples in panel A (marked by dash lines, mean±SD, n=5~6). One-way ANOVA test, *p<0.05, **p<0.01, and ***p<0.001. **C**. Fractional shortening (FS) of male zebrafish before (basal) and at different time points during recovery in DMSO or E2 (1 nM) after CI, as measured by echocardiography. One-way ANOVA with LSD post hoc test, *p<0.05, **p<0.01.

### Estrogen enhanced immune and inflammatory responses in regenerating heart

To elucidate the mechanism underlying the sexual difference in zebrafish heart regeneration, we performed comparative transcriptomic analyses on RNA extracted from female and male hearts at 7 dpc. A total of 1050 genes were found to be differentially expressed (by more than two-fold) between two sexes with statistical significance across 3 biological replicates (supplementary file 3). Gene ontology (GO) analyses revealed that most of the biological processes enriched for the female-biased genes are related to immunological processes, such as immune response, inflammatory response and chemotaxis for immune cells. On the other hand, male-biased genes are more diverse in function, ranging from protein homeostasis, stress response to muscle contraction (Fig. 5A). Gene set enrichment analysis (GSEA, Subramanian et al., 2005) confirmed that immune-related pathways are among the most sexually dimorphic in post-injured zebrafish hearts (Fig. S3). As female heart regeneration is faster, the over-representation of immune-related genes in heart-injured females might be a mere reflection of the fact that at 7 dpc the female hearts were at more advanced stage of regeneration than the males. A comparison of the published transcriptomic profiles of zebrafish hearts collected at 2 and 5 dpc (Lai et al., 2017), however, shows that immune-related genes were not among the most differentially expressed genes between these two days (Fig. S4), indicating that the observed female-biased expression of immune-related genes after heart injury was far more pronounced than can be explained by the difference in heart regeneration rate between sexes.

**Figure 5.**
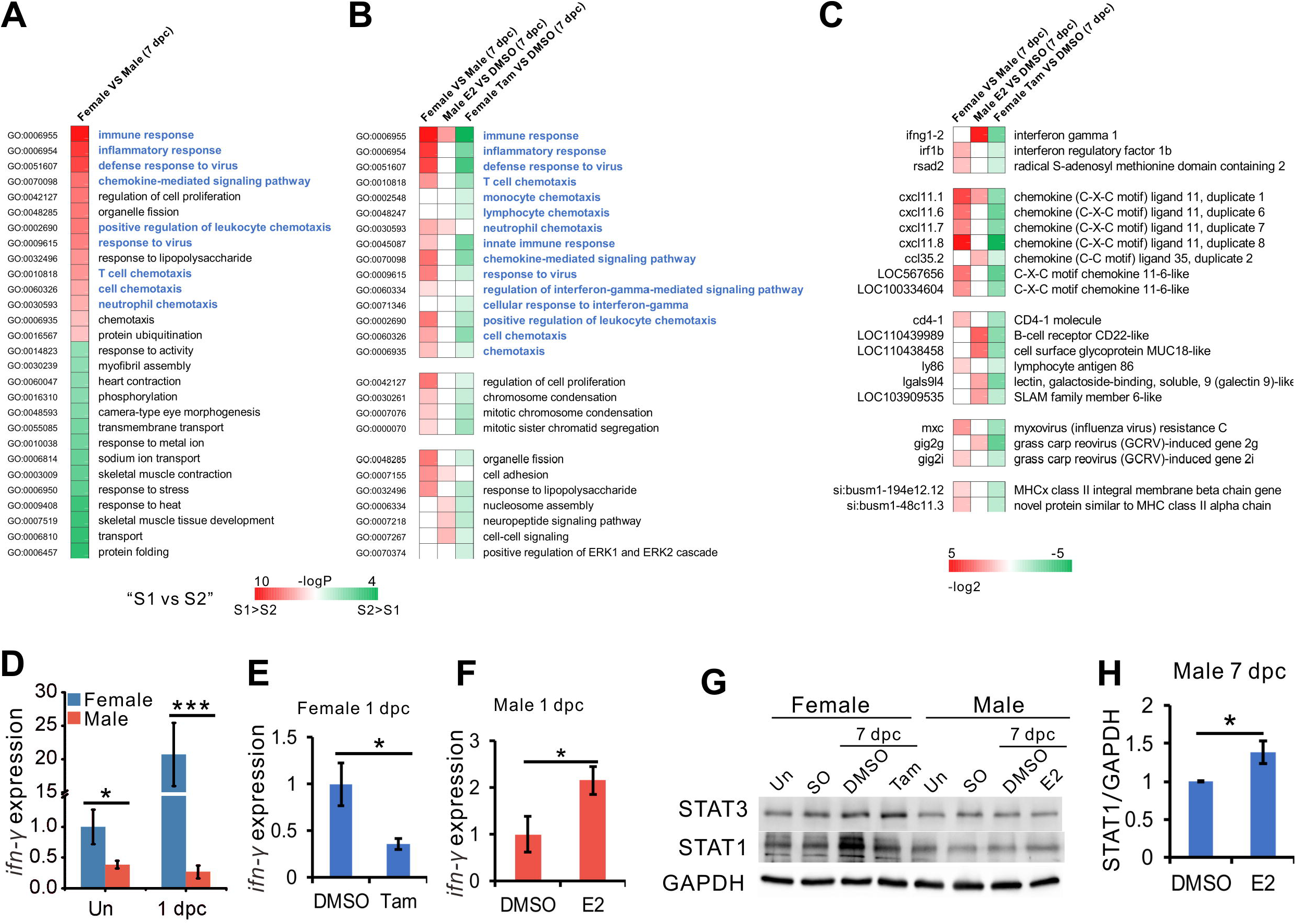
Estrogen induces inflammation in the injured zebrafish heart. **A.** Gene Ontogeny (GO) terms significantly enriched in genes differentially expressed in female vs male hearts at 7 dpc. **B**. Comparison of gene expression patterns in the hearts of female vs male, E2 (1 nM) -treated male vs untreated male, and Tamoxifen (1 μM) -treated female vs untreated female, all at 7 dpc. **C.** Expression of immune-nd inflammation-related genes in these datasets. **D-F**, Expression of *ifn-*γ in female and male heart of zebrafish at uninjured fish and 1dpc (D), tamoxifen decreased (E) and E2 increased (F) the expression of *ifn-*γ in female and male at 1 dpc. N=3, two tail t test, *p<0.05, **p<0.01, ***p<0.001. **G-H**, western blot of STAT1 and SATA3 in female and male heart at 7dpc after different treatment. H, bar chart to show the quantification of the expression of STAT1 in male fish in panel G, n=3, two tail t test, *p<0.05.

To test how much the observed female-specific gene expression pattern in heart regeneration was shaped by estrogen, the respective effects of E2 treatment of male fish, and tamoxifen treatment of female fish, on the transcriptome of injured hearts were investigated. Fig. 5B shows that tamoxifen treatment reversed almost all female-specific gene expression signatures in the injured heart, whereas genes involved in immune response and neutrophil chemotaxis were upregulated in estrogen-treated males. Among the genes whose expression level in injured hearts are sexually dimorphic, and/or reciprocally regulated by E2 and tamoxifen, are interferon-gamma (IFNg), interferon regulatory factor 1b (IRF1b) and various isoforms of CXCL11 (Fig. 5C).). In mammals, both IRF1 and CXCL11 are IFNg-inducible (Yang et al., 2007; Flodstro□m & Eizirik, 1997), suggesting the interferon-gamma pathway maybe instrumental to the sexual dimorphism in heart regeneration. Q-PCR analysis confirmed that the expression level of IFNg in the female heart increased by ~20-folds (1 dpc) after injury, while remained unchanged in males (Fig. 5D). IFNg expression after heart injury was significantly stimulated by E2 treatment of male fish and suppressed by tamoxifen treatment of females (Fig. 5E, F), in consistent with the role of estrogen in IFNg expression in other tissues (Fox et al., 1991; Hao et al., 2013).

In consistent with the female-specific induction of IFNg expression, we observed a significant increase of STAT1, a target of IFNg (Horvath, 2004), in regenerating female hearts but not in males (Fig. 5G), The upregulation of STAT1 in female hearts could be reversed by tamoxifen (Fig. 5G). In males, E2 treatment resulted in a small but significant increase in STAT1 (Fig. 5G, H). These results are consistent with the recent observations that STAT1 is an estrogen-responsive gene in mammals (Young et al., 2017). STAT3 is essential to injury-induced cardiomyocyte proliferation in general (Fang et al., 2013). Our data showed that STAT3 was also induced in female hearts after heart injury, but this induction was not tamoxifen-sensitive (Fig. 5G), suggesting STAT1 may be more important than STAT3 in orchestrating sexual differences in regeneration rates.

### Estrogen treatment preconditions zebrafish for faster heart regeneration

Although sham operation does not trigger heart regeneration, this process is known to precondition zebrafish for a more efficient regeneration if their hearts are damaged shortly afterwards, possibly due to an increase in inflammation as a result of thoracotomy (de Preux Charles et al., 2016a, b). We observed that sham operation led to significant increases in the expression of estrogen receptors (Fig. 2A, B) and plasma E2 (Fig. 2C, D) at 7 dpc. To test whether estrogen is instrumental to this preconditioning effect, we treated thoracotomised male zebrafish with estrogen for 2 days prior to cryoinjury (Fig. 6A). As expected, the regenerative programme, as judged by the number of proliferating cells and the expression of embCMHC, were enhanced by thoracotomy (Fig. 6B-D), confirming the occurrence of preconditioning effect. However, this effect was significantly reduced when the fish were exposed to tamoxifen during the preconditioning period (Fig. 6B-D). Furthermore, male zebrafish pretreated with E2 for 2 days prior to cryoinjury showed a significant increase in PCNA and embCMHC expression, when compared to DMSO pretreated fish. Hence, E2 pre-treatment could replace thoracotomy in inducing the preconditioning effect on heart regeneration via inflammation.

**Figure 6.**
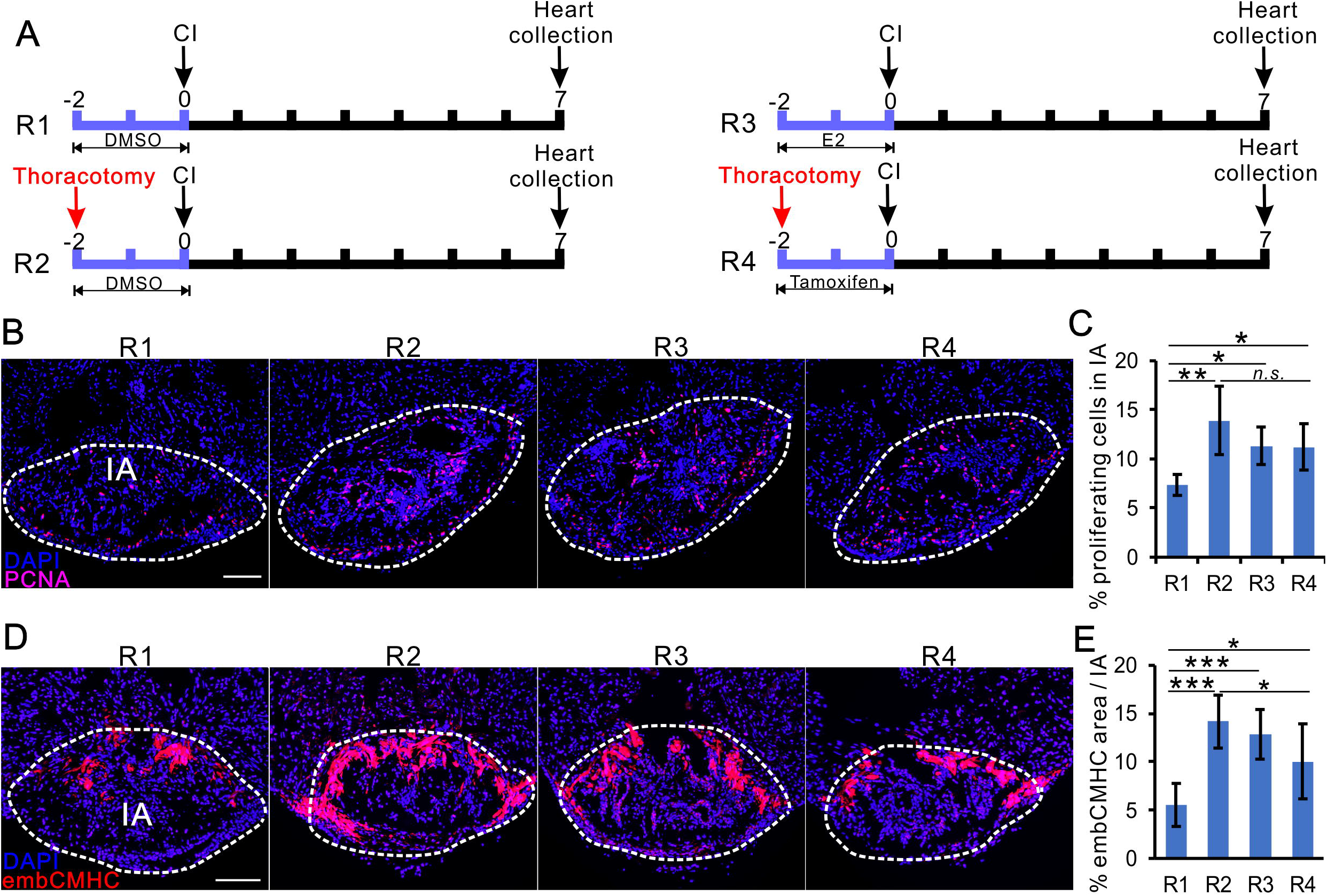
Estrogen precondition treatment promote zebrafish heart regenerative process. **A**. Design of the experiments that generated the data presented in this figure. **B**. PCNA immunofluorescence (red) in the male heart from regimens R1-R4. **C**. Quantification of PCNA-positive cells in panel B (mean±SD, n=5) in injured areas (marked by dash lines). Scale bars: 100μm. One-way ANOVA test, *p<0.05, **p<0.01, n.s. nonsignificant difference. **D**. embCMHC immunofluorescence (red) in the the male heart from regimens R1-R4. **E**. Quantification of embCMHC staining in panel E (mean±SD, n= 5~6) in injured areas. Scale bars: 100μm. One-way ANOVA with LSD post hoc test, *p=0.05, ***p<0.001.

## Discussion

In this study, we demonstrated by cellular, anatomical, physiological and biochemical (Fig. 1, 2) parameters that zebrafish heart regeneration is sexually dimorphic. As far as we know, this is the first evidence of sexual dimorphism in heart regeneration. Sexual differences in the regeneration of other tissues has been observed in mammals but has not been documented for the heart (Harada et al., 2003; Blankenhorn et al., 2003; Deasy et al., 2007). In zebrafish, the pectoral fin is the only tissue that has been shown to demonstrate a sexually dimorphic regenerative capacity (Nachtrab et al., 2011). Like their hearts, males regenerate the pectoral fin more slowly and often incompletely. However, this phenomenon was not observed in other fins (Nachtrab et al., 2011), suggesting that this sexual dimorphism is related to the unique role of pectoral fins in reproduction (Kang et al., 2013; McMillan et al., 2013).

We demonstrated the positive effect of estrogen on heart regeneration: E2 accelerated heart regeneration in males (Fig. 3, 4), and tamoxifen retarded it in females (Fig. 3, 4). Our gene expression analyses suggested that female hearts demonstrated a stronger immune and inflammatory responses to cryoinjury. In particular, the expression of IFNg, as well as IFNg-inducible factors, in regenerating hearts was highly sexually dimorphic and E2 sensitive (Fig. 5). As inflammation is essential for tissue regeneration (Eming et al., 2017), including cardiac repair (Frangogiannis, 2015), our observation is consistent with the higher heart regeneration rate in female zebrafish. Importantly, a comparison between fish species with different heart regeneration rates attributes the high heart regeneration capacity of zebrafish to its enhanced inflammatory response to heart injury (Lai et al., 2017). These authors also discovered that treatment with poly I:C, a viral mimic known to stimulate IFNg-responsive genes (Farina, et al., 2010), can enable heart regeneration in medaka, a species that is normally unable to repair heart injuries. The immune-promoting effects of estrogen have been well documented in fish and mammals (Burgos-Aceves et al., 2016; Taneja, 2018). In addition, sham-operation preconditioning induced inflammation and promote heart regenerative programme (de Preux Charles et al., 2016a, b), which could replace by E2 pre-treatment and block by tamoxifen treatment.

We have also revealed a previously unknown aspect of tissue regeneration in zebrafish: the spontaneous endocrine disruption after cardiac damage towards feminisation. Our data reveal that cardiac damage triggers the expression of estrogen receptors, notably *esr2a* (Fig. 2A, B) and secretion of E2 (Fig. 2C, D), in both sexes. Interestingly, a genetic variant of *esr2* has been identified as a risk factor of myocardial infraction (Domingues-Montanari et al., 2008). In addition, ERb, a mammalian homologue of *esr2a*, has recently been found to locate in mitochondria and play bioenergetic roles (Liao et al., 2015). We observed a ~9-fold increase in *esr2a* expression in female hearts at 7 dpc. Such dramatic increase in ER expression, compounded by the ~5-fold increase of plasma E2 level, will likely sensitise estrogen-responsive genes and amplify the estrogen-dependent inflammatory response of the female heart towards injury. We postulate that this can further enhance the female prominence in heart regeneration, and firmly establish the sexual dimorphism of this process (see Fig. 7 for a model that summarises the data presented in this study). The stimulation of estrogen level and receptor by heart injury also operates in male fish, hitherto to a lesser extent. Interestingly, the level of serum estrogen among acute myocardial infarction and unstable angina patients were found significantly higher than in the normal group and intensive care patients (Aksut et al., 1986), suggesting that cardiac lesions, but not other traumas, can lead to an increase in estrogen level in humans. We now show that the unique role of estrogen in heart repair is evolutionarily conserved. For example, endocrine disruption was not associated with zebrafish fin regeneration (this study) and estrogen (17α ethinylestradiol) does not affect the rate of fin regeneration in zebrafish larvae (Sun et al., 2019).

**Figure 7.**
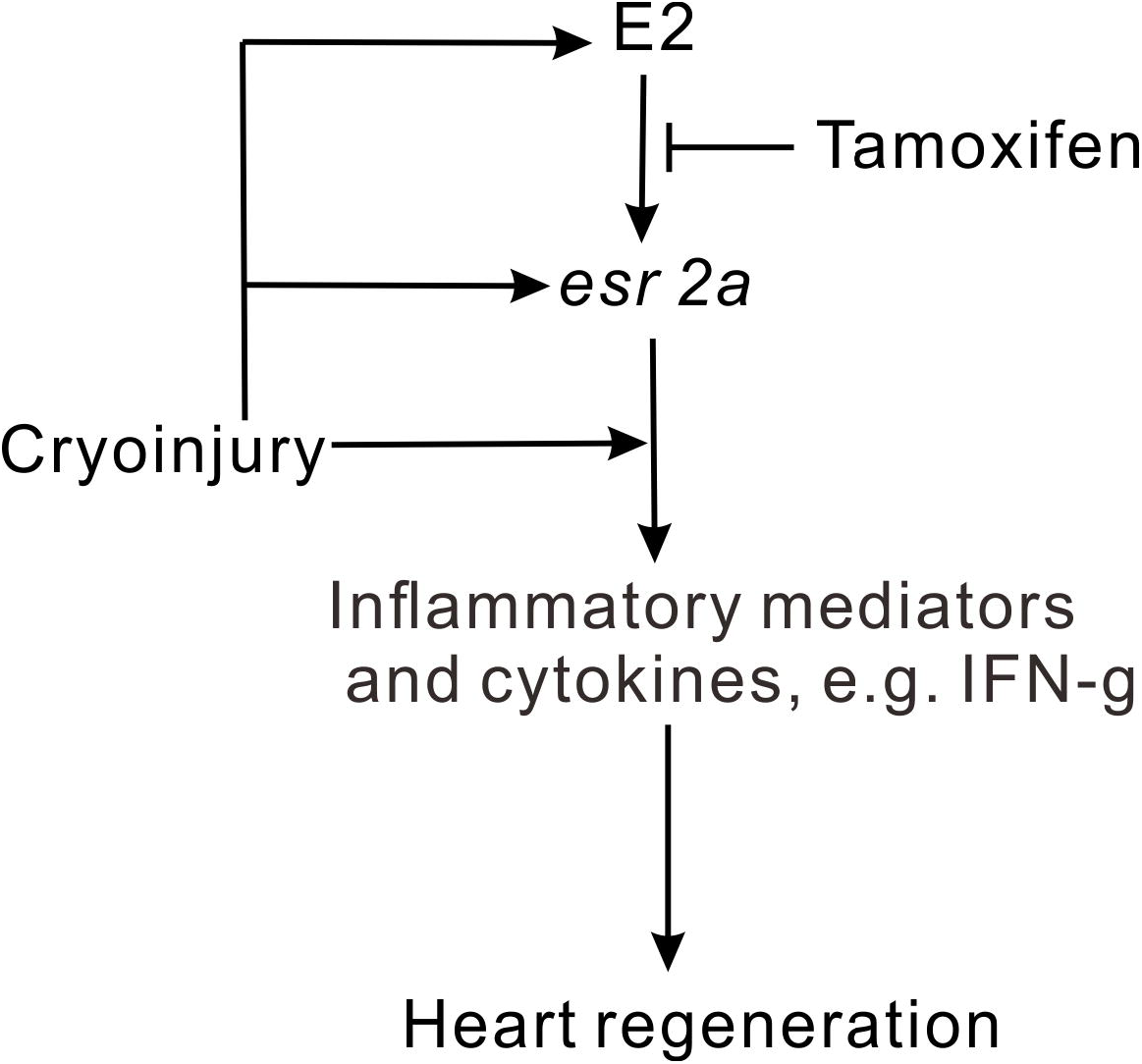
proposed model of role of estrogen in zebrafish heart regeneration. Cryoinjury of the heart triggers inflammation response and plasma estrogen upregulation of the fish. The increase of estrogen in plasma induced the expression of *esr2a*, which can be inhibited by tamoxifen. Estrogen promotes heart regeneration through *esr2a* induced and injury directly induced inflammation.

It is interesting to examine how the systemic increase of estrogen as a result of heart injury may impact the functions of other organs during heart regeneration. One of the consequences of the endocrine disruption is the detection of vitellogenin in males, an occurrence generally regarded as the hallmark of endocrine disruption by environmental agents (García-Reyero et al., 2004; Scott & Robinson, 2008). The accumulation of vitellogenin in regenerating hearts, but not in other tissues (Fig. 2, S2C), suggests a functional role in cardiac regeneration, rather than as a collateral consequence of estrogen secretion. The canonical function of vitellogenin is to transport lipids, calcium and phosphate to the developing oocytes (Arukwe & Goksøyr, 2003, Hara et al., 2016). We hypothesise that vitellogenin might play similar roles in the regenerating heart. Lipids are instrumental to cardiac remodelling in reptiles (Riquelme et al., 2011), and our previous micro-CT imaging of cardiac-damaged male zebrafish has shown the accumulation of lipids around the regenerating heart (Babaei et al., 2016). Future investigations will involve the characterisation of the vitellogenin cargoes isolated from the plasma and heart of fish after cardiac damage.

Like other major physiological challenges (Droujinine and Perrimon 2013; Kux and Pitsouli 2014), tissue regeneration likely requires the coordinated contributions from multiple tissue systems. This study reveals, for the first time, that the involvement of the endocrine and immune systems in zebrafish heart regeneration and opens up the use of zebrafish as an organismal model for the study of the sexual differences in human cardiac pathology.

## Materials and Methods

### Zebrafish maintenance

Zebrafish AB line was acquired from the Zebrafish International Resource Center (ZIRC; University of Oregon, Eugene, OR, USA). Fish were maintained in a recirculating system at 28°C with a photoperiod of 14 h of light/10 h of darkness, and fed with dry meal (TetraMin) three times a day, supplemented with live brine shrimp once a day (Westerfield, 2000). All the animal procedures used in this study were approved by the Department of Health, Hong Kong, SAR, China (refs (17-18) in DH/HA&P/8/2/5 Pt.1), and the experiments were conducted in accordance with the relevant guidelines and regulations in Hong Kong, SAR, China.

### Animal surgery

Adult zebrafish (12 to 18-month-old, the weight is 0.35-0.37g, with cardiac weight accounting for 0.5% of body weight) were anesthetised by immersion in system water containing 0.04% MS-222 (ethyl-3-aminobenzoate methanesulfonate salt; E10521; Sigma-Aldrich) for 3−5 min and then immobilized on a petri dish. After the removal of the ventral scales by forceps, a small incision was made through the body wall and the pericardium by using forceps and micro-dissection scissors, tearing the tissue rather than making a clean cut in order to facilitate healing. Once the pericardial sac was opened, the heart ventricle was exposed by gently squeezing the abdomen. For sham operation, the heart was gently pushed back into the body without any further treatment. For ventricular amputation, a portion of the ventricle was excised by using surgical fine scissors (Poss et al., 2002). For cryoinjury, the ventricle was touched for 10-12 seconds by metal probe pre-chilled in liquid nitrogen (Chablais et al., 2011). For fin amputation, the caudal fin of zebrafish was cut using a blade after anesthetization (Nachtrab et al., 2011). After the operation, fish were placed in a tank of fresh water, and reanimation was enhanced by pipetting water onto the gills for a couple of minutes. At time points indicated in the Results, the zebrafish were sacrificed by immersion in overdose concentration of MS-222. Hearts were dissected from the fish and their plasma were collected as previously described (Babaei et al., 2013).

### Chemical exposure

Adult male and female zebrafish, untreated or after surgery, were incubated in water containing 17β-estradiol (E2; E1132, Sigma-Aldrich) at 1 nM, tamoxifen (T9262, Sigma-Aldrich) at 1μM, or DMSO (D5879, Sigma-Aldrich), which was used to prepare the E2 and tamoxifen, at matching concentrations. The water was changed every day, and the fish were continuously exposed to the vehicle or drugs for durations indicated in Fig. 1–5.

### Western blotting

Proteins from zebrafish hearts were extracted by using RIPA Lysis and Extraction Buffer (89901, Thermo scientific) according to manufacturer’s instruction. BCA Protein Assay (23225; Thermo Scientific) was used to determine the total protein concentration of zebrafish plasma and heart protein extract, according to the manufacturer’s instruction. 20μg of protein sample per lane was separated in a 10% SDS polyacrylamide gel, transferred to polyvinylidene difluoride (PVDF) membrane (10600023; GE Healthcare life science) by using Mini-Trans Blot Electrophoretic Transfer Cell system (1703930; Bio-Rad Laboratories) according to the manufacturer’s instructions. The blot was blocked in 5% no fat milk in PBST (0.05% Tween 20 in 1× PBS) for 1 hour at room temperature, following by incubation in primary antibodies diluted in PBST overnight at 4 □. The following primary antibodies were used: mouse anti-GAPDH (60004-1; Proteintech Group, Inc.) at 1:10000, mouse anti-zebrafish vitellogenin, JE-2A6 at 1:2000, mouse anti-embCMHC at 1:500, rabbit anti-SATA1(R1408-2, Huabio Inc.) and rabbit anti-STAT3(ET1607-38, Huabio Inc) at 1:2000. For secondary antibodies, HRP-conjugated rabbit anti-mouse IgG (AP160P; Millipore) at 1:5000, and HRP-conjugated goat anti-rabbit IgG (AP307P; Millipore) at 1:5000 were used. The proteins were detected with the EMD Millipore Luminata Western HRP chemiluminescence substrate (WBLUF0500; Millipore) and the signals were visualized with the Western blotting system (C600; Azure Biosystems, Inc.). The band densities were quantified using Image J.

### Mass spectrometry

Proteins from plasma collected from individual fish (typically about 20 μg) were precipitated in 1 mL of acetone for 60 min at −20°C and pelleted at 1000g for 10 min at 4 °C. The pellet was air-dried (30 min) and then dissolved in 8M urea in 10 mM Tris-HCl (pH 8.0). After clarification (10000 g, 10 min, RT), the supernatant was reduced by 10mM dithiothreitol (R0861, Thermo Scientific) in 50 mM ammonium bicarbonate (09830, Sigma-Aldrich; 30 min, RT), and alkylated in 50mM iodoacetamide (90034, Thermo Scientific) in 50 mM ammonium bicarbonate (20 min, RT). LysC (Roche) was then added to the protein mixtures and incubated for 120 min at RT. The samples were then diluted with ammonium bicarbonate (50 mM, pH 8.0) so that the final concentration of urea was 1 M, followed by the addition of 1 μg trypsin (06369880103, Roche). The mixtures were incubated overnight at 37°C and dried in a vacuum centrifuge. Peptides were dissolved in a small amount of 0.1% trifluoroacetic acid (302031, Sigma-Aldrich; TFA), followed by purification with the use of ZipTip (MA01821, Millipore), following manufacturer’s instruction.

Mass spectrometry was performed on the Thermo Scientific Q-Exactive Plus Mass Spectrometer (Thermo Fisher Scientific, Bremen, Germany) coupled to a Thermo Ultimate 3000 RSLCnano HPLC system. About the 0.5 ug of the peptide mixture was loaded onto a 250nl OPTI-PAK trap (Optimize Technologies, Oregon City, OR) custom packed with Michrom Magic C18 5 μm solid phase (Michrom Bioresources, Auburn, CA). Chromatography was performed a Thermo EasySpray 25 cm × 75 μm C18 2 μm column, using 0.2 % formic acid in both Solution A (98%water/2% acetonitrile) and Solution B (80% acetonitrile/10% isopropanol/10% water), in a gradient from 2%B to 35%B over 140 min at a flowrate of 325 nL/min. The Q-Exactive Plus mass spectrometer was set up with a FT survey scan from 340-1500 m/z at resolution 70,000 (at 200m/z), followed by HCD MS/MS scans on the top 15 ions at resolution 17,500. The MS1 AGC target was set to 1e6 and the MS2 target was set to 2e5 with max ion inject times of 50ms and 75ms respectively. Dynamic exclusion placed selected ions on an exclusion list for 30 seconds. Charge exclusion was used to perform MS/MS only on +2, +3, and +4 ions.

MS/MS peak lists were exported as an .mgf file and proteins were identified via automated Mascot database searching (Matrix Science 2.2.04) of all tandem mass spectra against of the Danio rerio Protein Index protein sequence database that contained all zebrafish protein entries (143,725 sequences) from NCBI RefSeq (version 51). The instrument setting for the Mascot search was specified as “ESI-Trap.” Parameters used for the database search were as follows: a maximum of two missed cleavages; carbamidomethylation of cysteine as a fixed modification and oxidation of methionine, acetylation of protein N-term and Gln→pyro-Glu conversion as variable modifications; trypsin as the enzyme; a peptide mass tolerance of 10 ppm; a fragment mass tolerance of 0.6 Da; and an ion score of 35 as the cut-off, using a significance threshold of p < 0.05. After peptide identification, any peptides that had conflicting assignments were resolved, either one of identical proteins or by assignment to proteins with the largest number of peptides already present (by following Occam’s Razor principle).

The acquired MS data (in Thermo “.raw” format) from 3 biological replicates in the SO and Untreated groups (i.e., three separate plasma samples each collected from one individual fish) and 4 biological replicates in the VA group were quantitated by using Progenesis LC-MS (version 2.5, Nonlinear) as previously described (Babaei et al., 2013).

### Histology

Organs dissected from zebrafish were fixed with 4% paraformaldehyde at 4°C overnight, dehydrated and embedded in paraffin as previously described (Chablais et al., 2011). To measure scar size, the entire heart was serially sectioned at the thickness of 5μm. The sections were deparaffined, rehydrated and stained with picrosirus red (ab150681, Abcam). On each section, the labeled area and the area of the whole ventricle were measured by using ImageJ (National Institutes of Health, Bethesda, MA, USA), and the percentage of the scar volume to the entire ventricle was calculated by the summation of the data from all the sections (Xu et al., 2018). For the whole mount paraffin sections, the fish were fixed with 2% PFA and 0.05% Glutaraldehyde in 80% HistoChoice (H120, Amresco Inc.) with 1% sucrose and 1% CaCl_2_ at 4□ for 24h, and the paraffin sections were prepared according to Kong et al. (Kong et al., 2008).

For immunohistochemistry, antigen retrieval was performed on dewaxed sections in sodium citrate buffer (10 mM sodium citrate, 0.05% Tween 20, pH 6.0) at 95°C for 15 min. Primary antibodies used were: mouse anti-vimentin (ab8978; Abcam) at 1:200; mouse anti-PCNA (sc-56; Santa Cruz) at 1:200; mouse anti-zebrafish vitellogenin, JE-2A6 (V01408102; Biosense) at 1:200, mouse anti-embCMHC (N2.261;Developmental Studies Hybridoma Bank) at 1:50, rabbit polyclonal anti-GFP (ab13970; Abcam) at 1:200, mouse anti-L-plastin (sc-133218; Santa Cruz Biotechnology Inc.) at a dilution of 1:100. The secondary antibodies used were: Cy3-conjugated goat anti-mouse (A10521; Invitrogen) or Alexa Fluor 488-conjugated goat anti-rabbit (A11034; Invitrogen) antibodies at 1:500. The sections were amounted with cover slide in 50% Glycerol in PBS and images were acquired using Olympus BX61 microscope.

### Plasma E2 concentrations measurement

Plasma from three individual females or males were pooled, and three biological replicates were performed. The concentration of E2 in plasma were measured using Estradiol ELISA Kit (582251, Cayman).

### Quantitative PCR (q-PCR) and RNA sequencing

Total RNA was extracted from zebrafish heart, fin or liver using NucleoZOL reagent (740404; MACHEREY-NAGEL GmbH & Co. KG). Three individuals were pooled together as one biological replicate, and three biological replicates were performed. 1μg total RNA was decontaminated using RQ1 RNA-free DNase (M6101; Promega Co.) and then cDNA was synthesized using the PrimeScript reverse transcription (RT) reagent kit (6210B; Takara Bio. Inc.) according to the manufacturer’s instructions. The expression of each gene was determined by q-PCR using the SYBR Premix Ex Taq (RR402A; Takara Bio. Inc.), and *β-actin* was used as the reference gene. q-PCR was performed in triplicate for each gene, and the results were analyzed using the ΔΔ^CT^ method. All levels of expression of genes were normalized to *β-actin* and fold change was calculated by setting the normal female to 1. The primer sequences are list in Table 1.

**Table 1.**
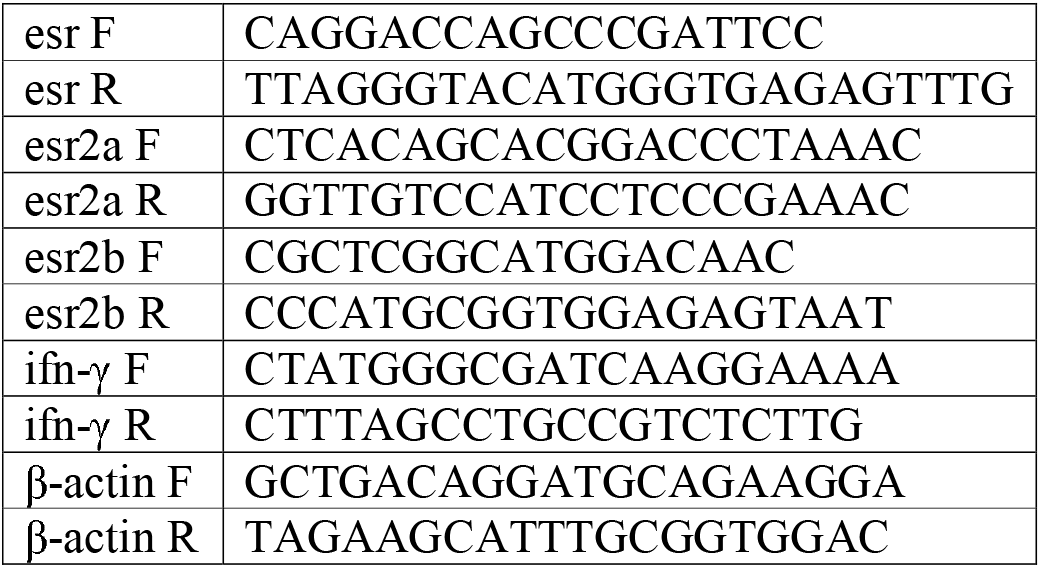
Primer sequence for Q-PCR

For RNA sequencing, total RNA was extracted and decontaminated, the sequencing was performed using BGISEQ-500 platform, averagely generating about 21.83 M reads per sample. The reference genome can be accessed at: http://www.ncbi.nlm.nih.gov/genome/50?genome_assembly_id=210873. The average mapping ratio with reference genome is 90.81%, the average mapping ratio with gene is 71.22%; A total of 25,900 genes were detected. The sequencing and the primary analysis were performed by BGI (Shenzhen, China), three bio-replicates of each samples were performed. We did comparative GSEA (Gene set enrichment analysis) and drew the union enrichment maps using an R package, HTSanalyzeR2 (https://github.com/CityUHK-CompBio/HTSanalyzeR2). Here, we focus on Biological Process Gene Ontology. By permutating 1000 times, we get the statistical significance and choose adjusted P value < 0.05 as significant results.

### Echocardiography

Zebrafish was anaesthetized with 0.02% MS-222 and fixed in a sponge upside-down. Echocardiographic studies were performed using Vevo LAZR Multi-modality Imaging Platform (FUJIFILM VisualSonics) under B-mode (50 MHz, 77 fps) and PW Doppler mode (40 MHz, 25 kHz PRF) at 20□ as previously described (González-Rosa et al., 2014; Hein et al., 2015; Wang et al., 2017). Ejection Fraction (EF) and Fractional Shortening (FS) were acquired using a plug-in of Vevo LAB (Vevo Strain) under B-mode images. The window was set to be 300 frames long, starting with diastalsis. PW Doppler images collected were analyzed using Vevo LAB (FUJIFILM VisualSonics) for E/A ratio calculation. 5 respective sets of E/A values were calculated for each sample.

### Quantification and Statistical Analysis

Three different images were taken for each heart. The percentages of proliferating cells in ventricle or injured area were the ratio of PCNA positive cells/DAPI. For untreated fish, all nuclei in the whole ventricle were counted; for cryoinjured fish, only the nuclei in the injured area (marked by white dash lines) in Fig. 1 and Fig. 6 were counted. The area of vimentin, embCMHC, and L-plastin expressions were quantified in the injured area using Image J and the percentage of the expression with respect to the area of the injured area was calculated (de Preux Charles et al., 2016a). For quantification of proliferating cardiomyocyte, the PNCA^+^/GPF^+^/ double positive cardiomyocyte within 100 μm of the vicinity of the injured area were quantified, and normalized with injured area in Fig. 3C. The data were expressed as the mean±S.D.(standard deviation). Statistical analysis was performed using Student’s two-tailed *t*-test and one way-ANOVA.

## Supplementary fiels

### Acknowledgements

SHC and YWL conceived and coordinated the study; SSX, SF and FB designed, performed and analyzed the experiments; FJX, KFW, and LS analyzed the echocardiographic data; TML and RR conducted the proteomic analyses; LNZ and XW do GSEA analysis; SSX and YWL wrote the paper; All authors reviewed the results and approved the final version of the manuscript. This work was supported by a general research fund grant [Project No. CityU 160213] from the Research Grants Council (RGC) of the Hong Kong Special Administrative Region, China, to SHC, and a Strategic Research Grant [Project No. CityU 7004661] from City University of Hong Kong to YWL. We thank the University Research Facility in Life Sciences of Hong Kong Polytechnic University for providing Vevo 2100 for this research. We thank all members of the SHC and YWL labs for fruitful discussions.

### Competing Interests

The authors declare that they have no conflicts of interest with the contents of this article.

---

**Figure S1. L-plastin positive leukocytes is sexually dimorphic in cryoinjured zebrafish heart. A**. L-plastin labeled leukocytes in the injured area at 1 dpc in female and male fish. **B**. Quantification of L-plastin positive cells in panel A, N=5, two tail t-test, *p<0.05. Scale bars: 100μm.

**Figure S2. Vitellogenin accumulates in the male zebrafish heart after cardiac damage. A**. Ratio of the relative abundance of selected plasma proteins on days 1, 3 and 7 after surgery. VA: ventricular amputation, SO: sham operation. N=3-5. **B**. Relative abundance of known acute response proteins in zebrafish plasma collected on day 1 (D1), day 3 (D3) and day 7 (D7) after ventricular amputation (VA) and sham operation (SO). Colours represent the log2 ratio of the abundance of each protein in VA vs SO. N=3-5, only data with p<0.06 are shown. **C**. Vitellogenin (VTG) immunofluorescence (green) in male zebrafish heart, gill, kidney and liver on day 5 after VA. Scale bars: 200um.

**Figure S3. Comparative transcriptomic analyses between female and male zebrafish after 7 days cardiac injury. A**. Enrichment map to show the gene profile in female and male heart at 7 dpc. Red color represents NES>0, and geneset upregulated; blue color represent NES< 0, and geneset downregulated. The strength of the red or blue color represent the value of the adjusted p-value. **B**. GSEA plot to show the location of the maximum enrichment score (ES) and the leading-edge subset of immune response gene.

**Figure S4. Gene Ontogeny enrichment analysis on the differentially expressed genes in zebrafish hearts at 2 vs 5 dpc**. Based on data published in Lai et al., 2017.

