## Supplementary figures and images for "Estrogen accelerates heart regeneration by promoting inflammatory responses in zebrafish"

### Fig S3

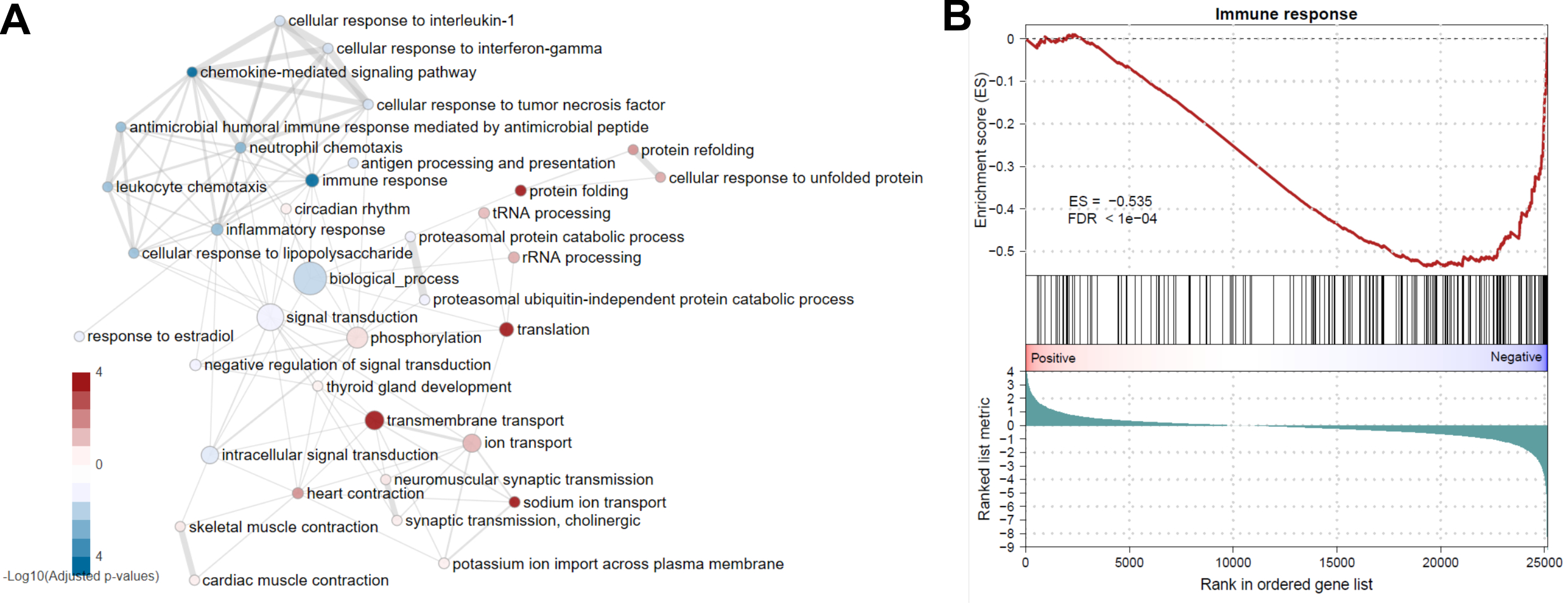

### Fig S4

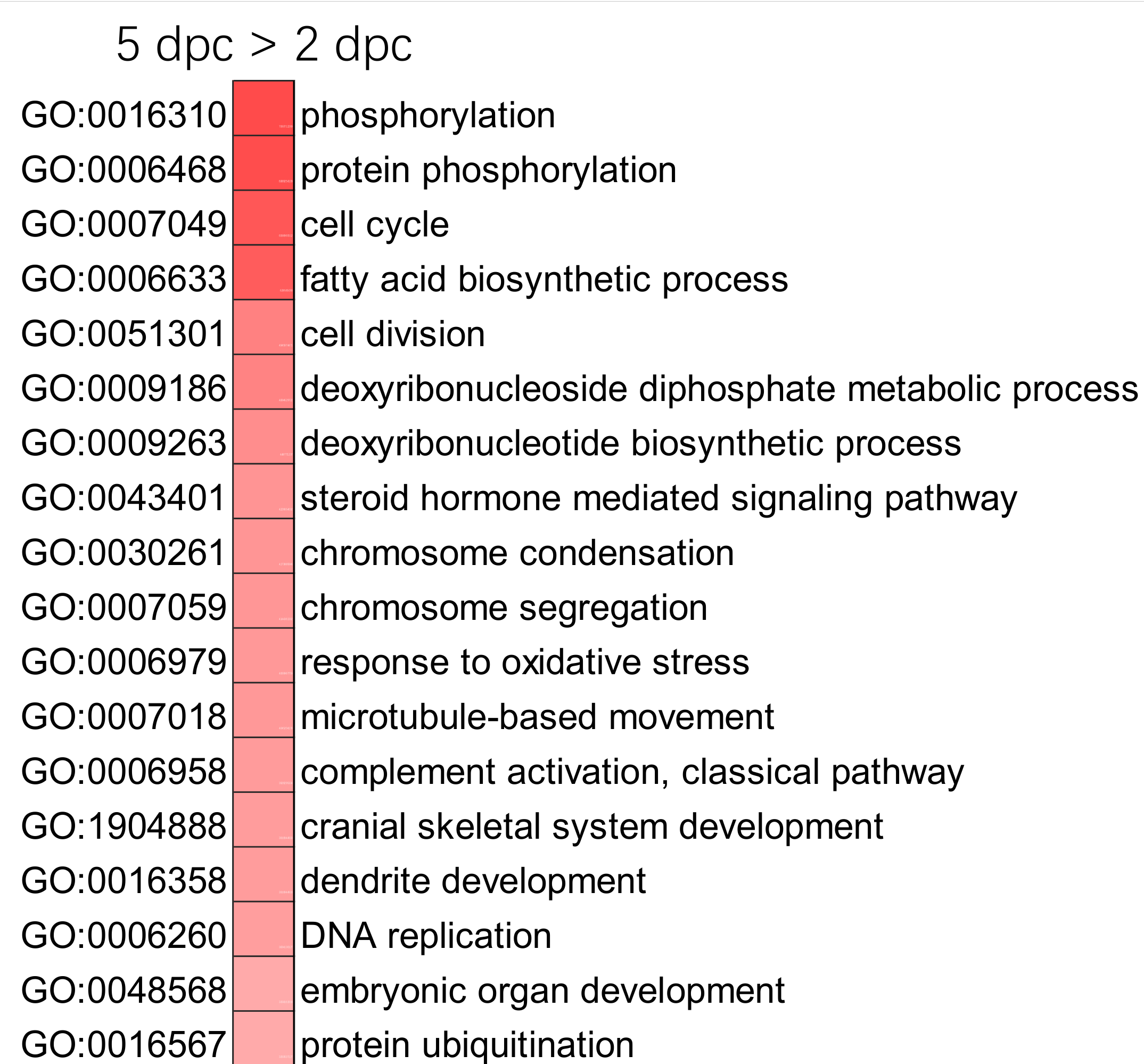

### Fig. S1

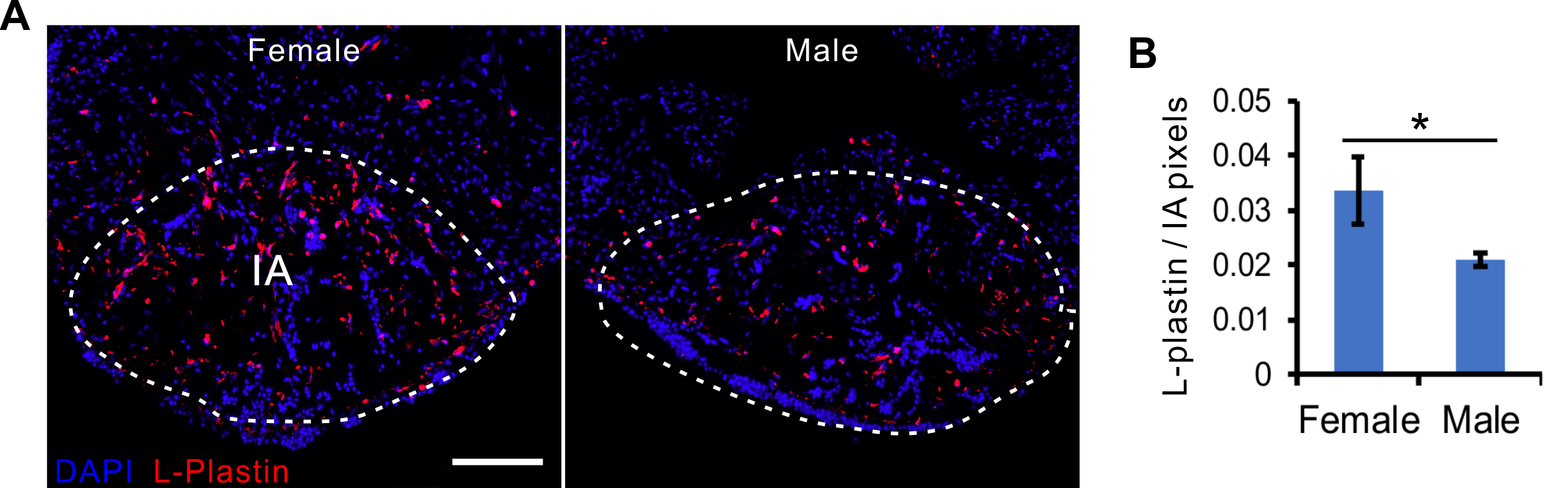

### Fig. S2

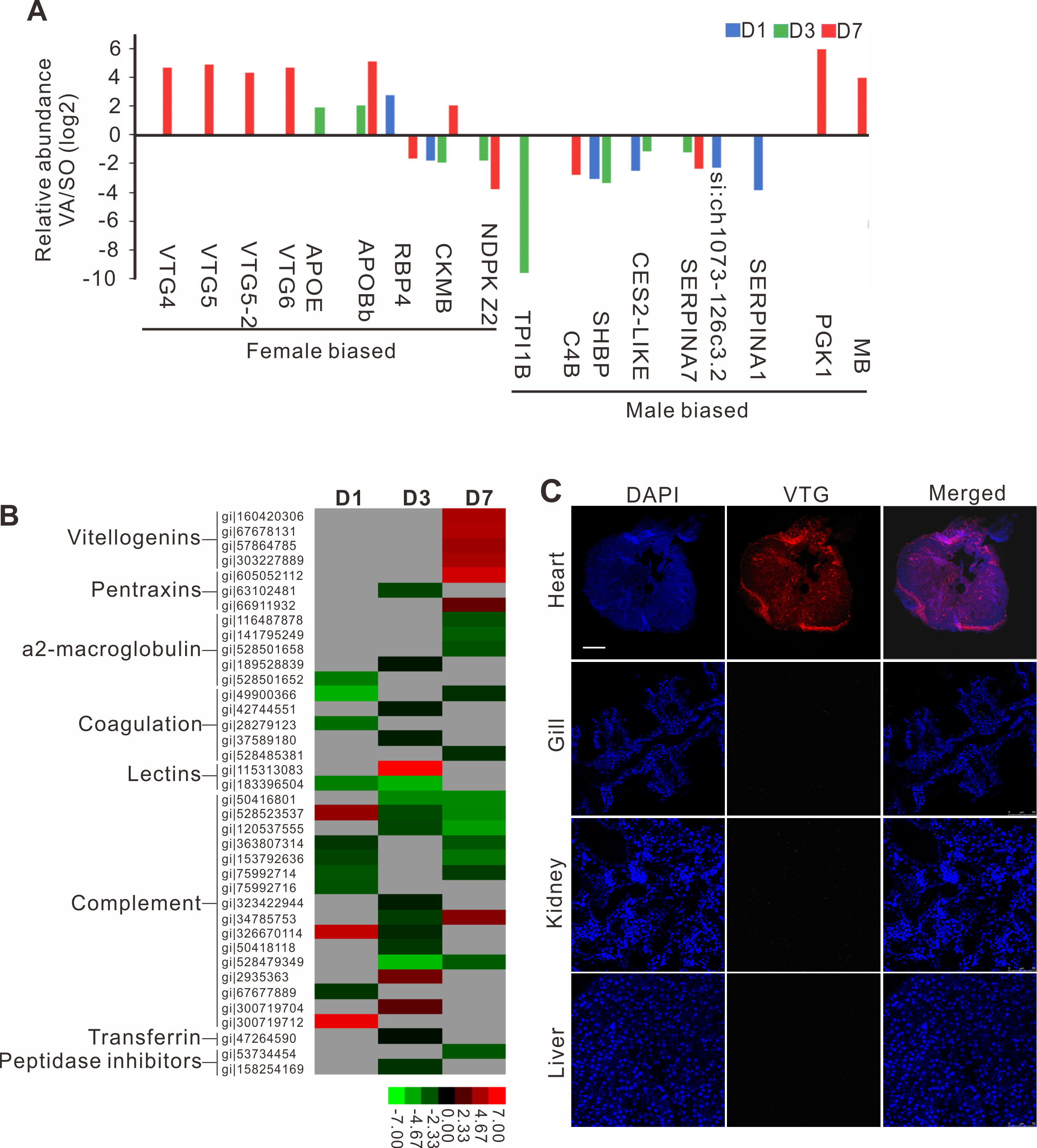
